# External light dark cycle shapes gut microbiota through intrinsically photosensitive retinal ganglion cells

**DOI:** 10.1101/2020.10.23.351650

**Authors:** Chi-Chan Lee, Tsung-Hao Lu, I-Chi Lee, Yan-Fang Zou, Hon-Tsen Yu, Shih-Kuo Chen (Alen)

## Abstract

Gut microbiota has been shown to involve in many physiological functions such as metabolism, brain development, and neuron degeneration disease. Intriguingly, many microbes in the digestive tract do not maintain a constant level of their relative abundance but show daily oscillations under normal conditions. Recent evidence indicates that chronic jetlag, constant darkness, or deletion of the circadian core gene can alter the composition of gut microbiota and dampen the daily oscillation of gut microbes. These studies suggest that the interaction between the host circadian clock and the light-dark cycle plays an important role in gut homeostasis and microbiota. However, how or whether environmental factors such as the light-dark cycle could modulate gut microbiota is still poorly understood. Using genetic mouse models and 16s rRNA metagenomic analysis, we found that light-dark cycle information transmitted by the ipRGC-sympathetic circuit was essential for daily oscillations of gut microbes under temporal restricted high fat diet condition. Furthermore, aberrant light exposure such as dim light at night (dLAN), acting through intrinsically photosensitive retinal ganglion cells (ipRGCs), could alter the composition, relative abundance, and daily oscillations of gut microbiota. Together, our results indicate that external stimulation, such as light-dark cycle information, through the sensory system can modulate gut microbiota in the direction from the brain to the gut.

## Introduction

Gut microbiota could influence the central nervous system through a proposed gut-brain axis to modulate neuronal development, function, and degeneration. It has been shown that germ-free mice have different hypothalamus metabolites compared to normal mice. In addition, germ-free or antibiotic-treated mice have different gene expression profiles in the CNS and lower anxiety levels compared to normal SPF mice(1–4). The development of the endocrine neuron such as oxytocin neurons is affected by the gut microbe during the postnatal period, which will influence the adult stage social interaction(5). Finally, the gut microbiota will also influence the aggregation of alpha-synuclein in the gut, which could lead to Parkinson’s disease(6). The composition of gut microbiota and the relative abundance of specific microbes could be influenced by the host’s genetic factors or feeding scheme. However, the relative abundance of many gut microbes displays daily oscillation even under normal condition(7–10). Although the direct physiological implication of the gut microbe daily oscillation is poorly understood, a recent report showed that arrhythmic gut microbe is associated with patients with type 2 diabetes(11). Recent evidence showed that chronic jetlag, constant darkness, and reversal of the light-dark cycle could disrupt the daily oscillation of gut microbe (9, 10, 12, 13). However, the mechanism for how light dark cycle could influence the daily oscillation of gut microbes is unclear. In addition, mice with clock gene knockout also display arrhythmic gut microbiota(10, 13). These studies suggest that both external and internal factors could influence the daily oscillation of gut microbe, and their interaction may play an important role for gut homeostasis and microbiota (9, 10, 12, 14–16). However, how do external factors influence the gut microbiota is unknown.

One candidate for modulation of daily gut microbe oscillation is the circadian clock system. This system is entrained to the daily light-dark cycle (LD) via input from the melanopsin-expressing, intrinsically photosensitive retinal ganglion cells (ipRGCs) to the suprachiasmatic nucleus (SCN) (17–22). In addition, ipRGCs also influence many other non-image-forming functions by projecting to brain regions in the thalamus and hypothalamus such as the olivary pretectal nucleus for pupillary light reflex (23–26). Genetic elimination of ipRGCs using the Opn4-DTA or Opn4-Cre; DTR mouse lines impairs the ability of mice to transmit external light-dark cycle information for circadian photoentrainment which causes these mice to “free run” under any kind of environmental light-dark cycle (17, 27, 28). On the other hand, knockout of the photopigment melanopsin (MKO) only produced a light detection phenotype for non-image-forming functions under high light intensity (18, 26, 29), which suggests that ipRGCs transmit signals to the brain by combining high luminance signals detected via melanopsin and low luminance signals originating from the canonical rod and cone photoreceptors. Recent evidence showed that ipRGCs can modulate many additional physiological functions such as emotion, hair regeneration, and body temperature (30–32). Since ipRGCs provide environmental luminance signals for many non-image-forming visual functions, it is likely that they could also influence gut microbiota. Using 16s rRNA analysis and various kinds of ipRGC-related mutant mice, here we show that light-dark cycle information could drive the daily oscillation of gut microbes through the ipRGCs-sympathetic nerve circuit. Furthermore, aberrant light-dark cycles such as light exposure during the night time could cause dysbiosis and dampen the daily oscillation of gut microbes.

## Results

### Light/dark cycle information is important to drive daily oscillations of gut microbes

To determine how the light-dark cycle may directly influence the composition of gut microbiota, we designed a specific dim light at night (dLAN) conditions to provide light exposure while minimizing the disruption of the central circadian clock. Furthermore, mice were under temporal restricted feeding cycle with high fat diet to eliminate daytime feeding behavior which disrupts circadian clock (33) (supplementary figure 1 and supplementary information). Under the dLAN condition, we could test the influence of additional light exposure during the nighttime while keeping mice entrained to 24 hours daily light-dark cycle. We first compared the gut microbiota under normal LD condition, under constant darkness (DD) with reduced light exposure, or under dLAN condition with extra light exposure. After 2 weeks of normal LD cycle, constant darkness, or dLAN conditions, fecal samples were collected from WT mice at six time points throughout the day. We perform 16S rRNA based next-generation sequencing and use the operational taxonomic unit (OTU) based JTK_CYCLE (34) to analyze the daily gut microbial oscillation in mice housed under different light-dark cycles. For WT mice, many microbes display daily oscillations under LD conditions (Fig. 1A, C, and D). Surprisingly, although the locomotor activity of mice and most circadian clock genes were entrained to 24 hours cycle under dLAN condition (Supplemental Figure 1), percentages of oscillating gut microbes were highly reduced in mice housed under dLAN conditions (Fig. 1A, C, and D). We also confirm that the daily oscillation of gut microbes is significantly reduced in mice housed under the DD condition, even with free running endogenous clock (Fig. 1B, C, and D). The temporal restriction feeding with high fat diet is not sufficient to drive the full daily oscillation of gut microbe for mice housed under dLAN and constant darkness conditions. Together, our results suggest that the environmental light-dark cycle is one of the major factors driving the daily oscillation of gut microbe.

**Figure 1.**
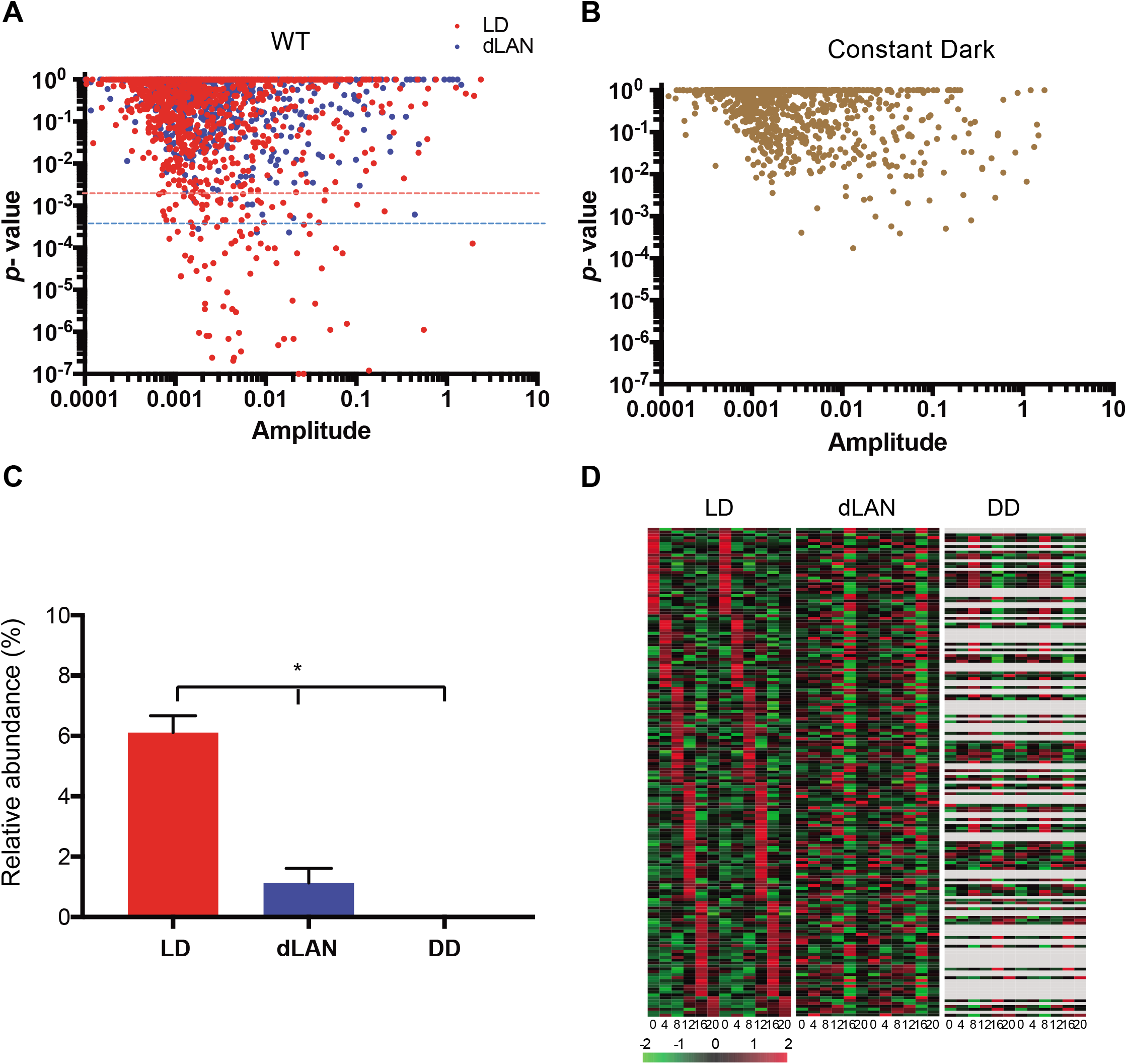
Light-dark cycle is important to drive the daily oscillation of gut microbes. (**A-B**) OTU based JTK_CYCLE analysis shows diurnal oscillating microbes from WT mice housed under normal light-dark cycle (A, LD), dim light at night (A, dLAN), and constant darkness (B) conditions. Dash lines indicate the Benjamini-Hochberg q-value < 0.05. (**C**) Summation of relative abundance from all oscillating OTUs in WT mice housed under LD, dLAN and constant darkness (DD) conditions. (**D**) Relative abundance heat map of oscillating OTUs detected by the JTK_CYCLE. Each row represents 1 OTU across the day and the graph is double plotted. Gray color indicates OTU not found. n = 4-5 for each time point. Graphs without dash line indicate that the q-values for all OTU were higher than 0.05.

### Daily oscillation of gut microbes is driven by light-dark cycle through ipRGCs

It has been shown that ipRGC is the primary conduit to transmit light information for many non-image-forming visual functions such as circadian photoentrainment and pupillary light reflex in mammals. Although the free running central clock could not fully drive the daily oscillation of gut microbes, ipRGCs and/or melanopsin signaling may be involved in transmitting light-dark cycle information to shape the gut microbiota. To test this hypothesis, we analyzed daily oscillation of gut microbe using JTK_CYCLE from control, MKO, and ipRGC eliminated (Opn4^DTA/DTA^) mice housed under normal LD cycle or dLAN condition (Fig. 2A-C). First, we compared the gut microbe oscillation between control and MKO mice. Interestingly, the pattern of daily oscillations in MKO differed from that in control mice under LD condition, while the total oscillation percentage in MKO is still similar to control (Fig. 2D and E). In control mice, peaks of oscillating OTUs were spread relatively evenly throughout the day (Fig. 2F), whereas in MKO mice, peak times for many oscillating OTUs occurred between ZT8 - ZT12 (Fig. 2G). These *de novo* microbe oscillations in MKO mice suggested that rod and/or cone signals might partially compensate for the loss of melanopsin. Next, we test whether ipRGC is the sole conduit to transmit melanopsin and rod/cone signal for gut microbe daily oscillation by comparing the rhythmicity between control and Opn4^DTA/DTA^ mice. Because the running activities of ipRGC eliminated (Opn4^DTA/DTA^) mice were not tied to the environmental light-dark cycle, JTK_CYCLE analysis for Opn4^DTA/DTA^ mice was performed by arranging data points to match either circadian time (CT) according to the wheel running activity (Supplementary Fig. 2) or zeitgeber time (ZT) according to the light-dark cycle. Strikingly, in Opn4^DTA/DTA^ mice, daily gut microbial oscillations were greatly attenuated (< 3%) under both LD and dLAN conditions when analyzed with either CT or ZT time (Fig. 2H). This result indicates that ipRGC is essential to provide normal light-dark cycle information from melanopsin and rod/cone to drive the gut microbe daily oscillation.

**Figure 2.**
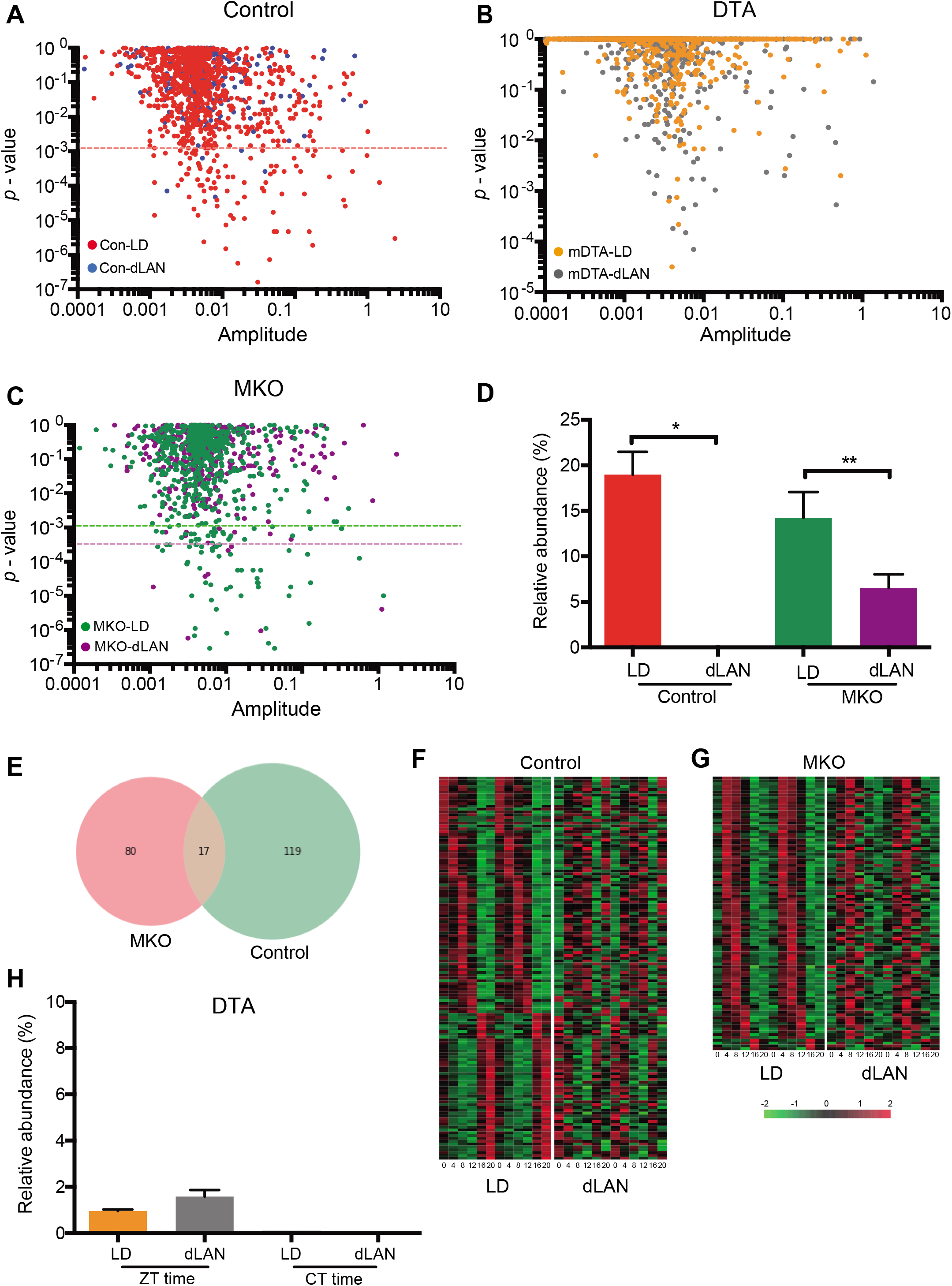
Light-dark cycle information transmitted by ipRGC is required for gut microbe oscillation. (**A-C**) Relative abundance heat map of oscillating OTUs detected by the JTK_CYCLE (p < 0.05, Benjamini-Hochberg q-value < 0.05) in control (A), Opn4^DTA/DTA^ (B), and MKO (C) mice housed under LD and dLAN conditions. (**D**) Summation of relative abundance from all oscillating OTUs in control and MKO mice. (**E**) Vann diagram of oscillating OTUs in control and MKO mice. (**F-G**) Relative abundance heat map of oscillating OTUs detected by the JTK_CYCLE in LD or dLAN conditions from control (F) or MKO (G) mice. Each row represents 1 OTU across the day and the graph is double plotted. (**H**) Summation of relative abundance from all oscillating OTUs in Opn4^DTA/DTA^ mice according to the circadian time (CT) of the mice, or the light/dark and food cycle (ZT). n = 4-5. * p<0.05, ** p < 0.01. All bar graphs are presented as mean ± SEM.

### Light signals transmitted by ipRGC influenced gut microbiota composition

In addition to modulate the daily oscillation of particular gut microbes, we next asked whether ipRGCs could modulate the overall composition of gut microbiota. We used principal coordinate analysis (PCoA) to compare the gut microbiota composition between control, MKO, and Opn4^DTA/DTA^ mice. Analysis of mean and variation, using ANOSIM (35) and Adonis (PERMANOVA) (36) respectively, showed that gut microbiota composition is significantly different between control and MKO mice (Fig. 3A), and also significantly different between control and Opn4^DTA/DTA^ mice (Fig. 3B). The alpha diversity (richness) was significantly lower for MKO and Opn4^DTA/DTA^ mice compared to control mice (Fig. 3C); while beta diversity (variation between samples) from MKO and Opn4^DTA/DTA^ mice were both significantly higher than control mice (Fig. 3D). These results indicate that melanopsin signaling and ipRGCs are important factors maintaining normal gut microbiota composition. Next, we compared the gut microbiota from control, MKO, and Opn4^DTA/DTA^ mice housed under LD or dLAN condition. The PCoA analysis showed clear separation of gut microbiota from control mice housed under LD versus dLAN conditions (Fig. 3E), indicating night time light exposure could modulate overall gut microbiota in mice. In contrast, neither MKO nor DTA animals showed separation of gut microbiota when mice housed in LD or dLAN condition (Fig. 3F and G). Moreover, neither alpha nor beta diversity of microbiota differed between LD and dLAN conditions in MKO and Opn4^DTA/DTA^ mice, though they were significantly different in controls (Fig. 3C-D). Since Opn4^DTA/DTA^ free run under any light-dark cycle, the activity time of mice was sometimes synchronized to the feeding schedule and then desynchronized to the feeding schedule during the course of the experiment. Interestingly, PCoA analysis resulted in a strong separation of gut microbiota between Opn4^DTA/DTA^ mice with their circadian clock synchronized or not synchronized to the feeding schedule under any light-dark cycle (Fig. 3H). These results showed that dyssynchronization between the feeding schedule and the endogenous circadian clock could modulate the gut microbiota in ipRGC eliminated mice. However, aberrant light-dark cycle induced dysbiosis is blocked in both melanopsin knockout or ipRGC eliminated mice.

**Figure 3.**
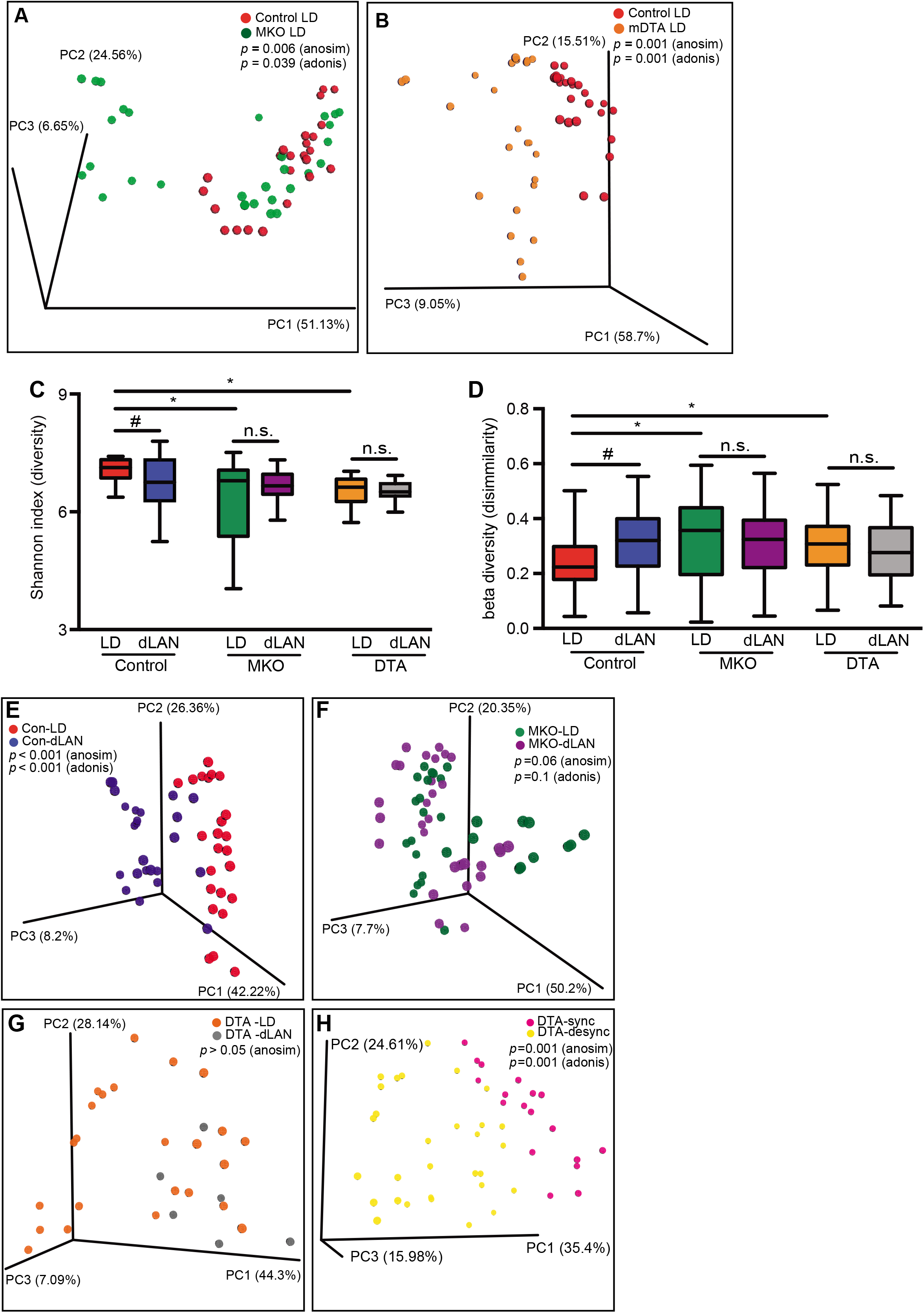
dLAN altered composition of gut microbiota. (**A**) Principal coordinate analysis (PCoA) of gut microbiota using weighted UniFrac matrix from control and MKO mice housed under LD conditions. (**B**) PCoA of gut microbiota from control and Opn4^DTA/DTA^ mice housed under LD conditions. The gut microbe compositions are significantly different between control and ipRGC manipulated mutant mice with both ANOSIM and Adonis analysis. (**C**) Box plot of the Shannon index represents alpha diversity for gut microbiota from control, MKO and Opn4^DTA/DTA^ mice under LD and dLAN condition. (**D**) Box plot for beta diversity generated from the weighted UniFrac distance matrix from control, MKO and Opn4^DTA/DTA^ mice under LD and dLAN condition. (**E-G**) PCoA of gut microbiota from control, MKO and Opn4^DTA/DTA^ (with similar phase angle difference to the light-dark cycle) mice housed under LD or dLAN conditions. Both ANOSIM and Adonis had significant differences for control mice housed under LD or dLAN conditions (E), but not for MKO (F) and Opn4^DTA/DTA^ mice (G). (**H**) PCoA of gut microbiota from Opn4^DTA/DTA^ mice either synchronized or desynchronized to the light/dark and feeding cycle. For box plots (C and D), error bars indicate minimum to maximum. The line represents the mean between groups; * p < 0.01; # p < 0.05 between same genotype using student t-test; n.s. not significant.

Next, we compared the relative abundance of each gut microbe under different light-dark cycles. The apparent gut microbe composition at the phylum level did not differ significantly among groups (Fig. 4A). However, linear discriminant analysis at multiple phylogenetic levels showed that thirteen classifications, including the genus *Rikenella*, *Odoribacter*, and *Lactobacillus*, were significantly different in control mice housed under dLAN versus LD conditions (Fig. 4B), whereas in MKO, only one classification had significant differences between dLAN and LD conditions (Fig. 4C). Strikingly, there were 23 classifications with significant differences between dLAN and LD conditions in Opn4^DTA/DTA^ mice. However, none of them were included in the 13 differential classifications in control mice housed under LD and dLAN conditions (Fig. 4D). Since Opn4^DTA/DTA^ mice free-run under our 12h: 12h light-dark cycle, the difference of microbiota pattern from Opn4^DTA/DTA^ mice may, again, be caused by dyssynchronization between the feeding schedule and the endogenous circadian clock. Together, our results suggested that light information through activation of melanopsin and/or transmitted by ipRGCs could influence the composition and diversity of gut microbiota.

**Figure 4.**
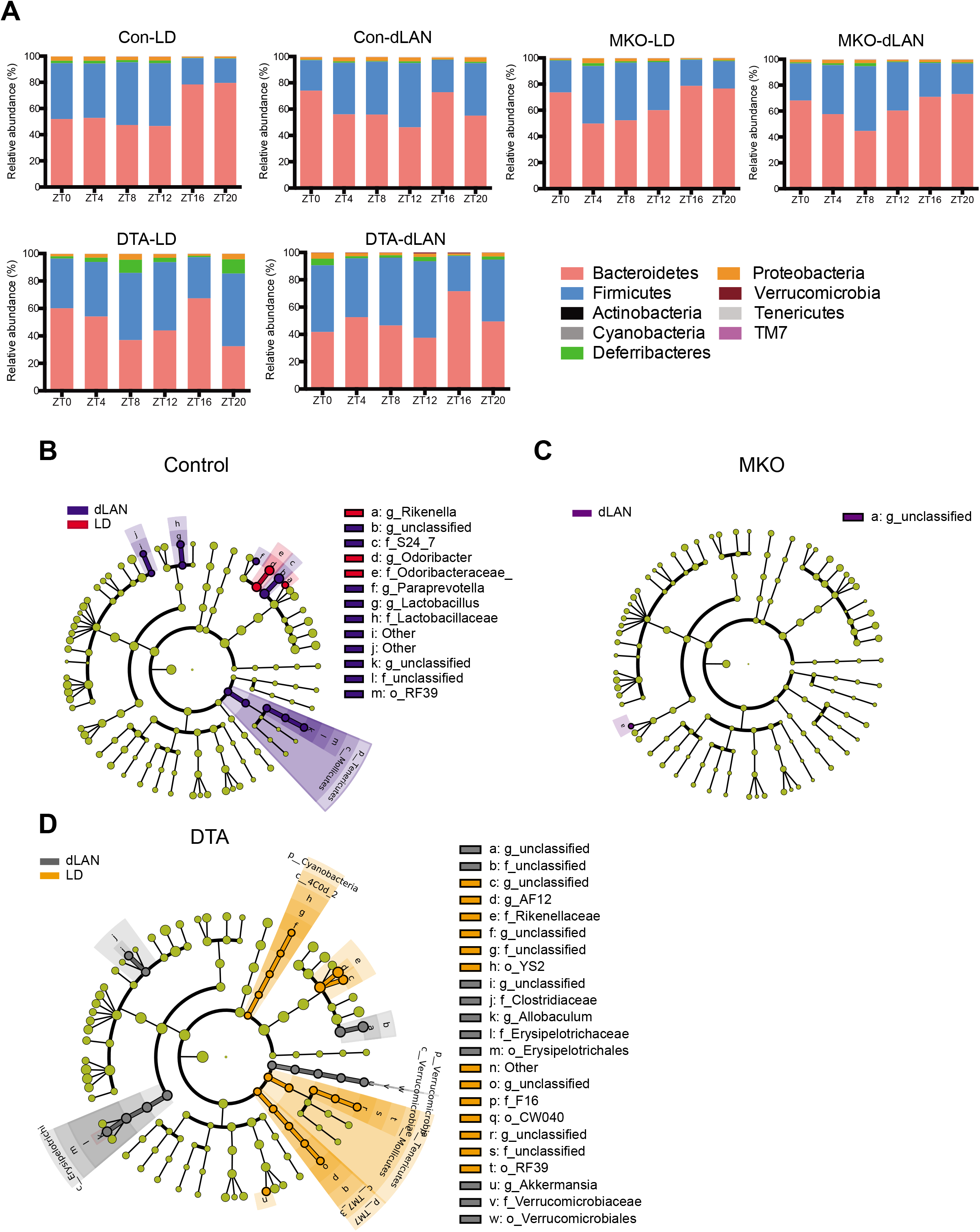
Enrichment of specific microbes from mice housed under dLAN and LD conditions. (**A**) Mean relative abundance of phylum from control (Con), melanopsin knockout (MKO) and ipRGC eliminated (Opn4^DTA/DTA^) mice housed under conditions of normal light/dark (LD) or dim light at night (dLAN). (n = 4-5 from each group) (**B**) The Cladogram shows the difference in relative abundance of microbes from control mice housed under conditions of LD and dLAN. Microbes in red have significantly higher relative abundance in LD conditions. Microbes in blue have significantly higher relative abundance in dLAN conditions. (**C**) The Cladogram shows the difference in relative abundance of microbes from MKO mice under LD and dLAN conditions. (**D**) The Cladogram shows the difference in relative abundance of microbes from Opn4^DTA/DTA^ mice under LD and dLAN conditions. For RF39, it is higher in the dLAN condition from control mice but higher in the LD condition from Opn4^DTA/DTA^ mice.

### ipRGC drive light-evoked, daily oscillations of gut microbes via sympathetic nerves

It has been shown that light can influence peripheral organs via the retinohypothalamic tract, independent of the circadian clock through autonomic circuits (37). Therefore, to determine the neural circuit responsible for the light-evoked modulation of gut microbiota, we tested whether the sympathetic nerve was involved in dLAN-induced dysbiosis. To eliminate sympathetic nerve function, WT mice were treated with 6-Hydroxydopamine (6-OHDA) (38). Body weight of mice decreased during the first week of 6-OHDA injection (Fig. 5A), consistent with the successful elimination of sympathetic nerve function (39). After body weight stabilized, mice were housed under either dLAN or LD conditions for 2 weeks. Sympathetic nerve ablation significantly dampened the daily oscillations of gut microbes under both LD and dLAN conditions (Fig. 5B and C), similar to our observations in ipRGC-ablated animals. These data suggest that light input driving daily oscillations occurs via the sympathetic nervous system. The dLAN induced beta diversity up-regulation is also blocked in sympathetic nerve ablation mice (Fig. 5D). Interestingly, sympathetic nerve ablation failed to abolish the effects of dLAN induced dysbiosis, as dLAN still altered the composition of gut microbiota under this manipulation (Fig. 5E), suggesting that these influences occur via a distinct circuit.

**Figure 5.**
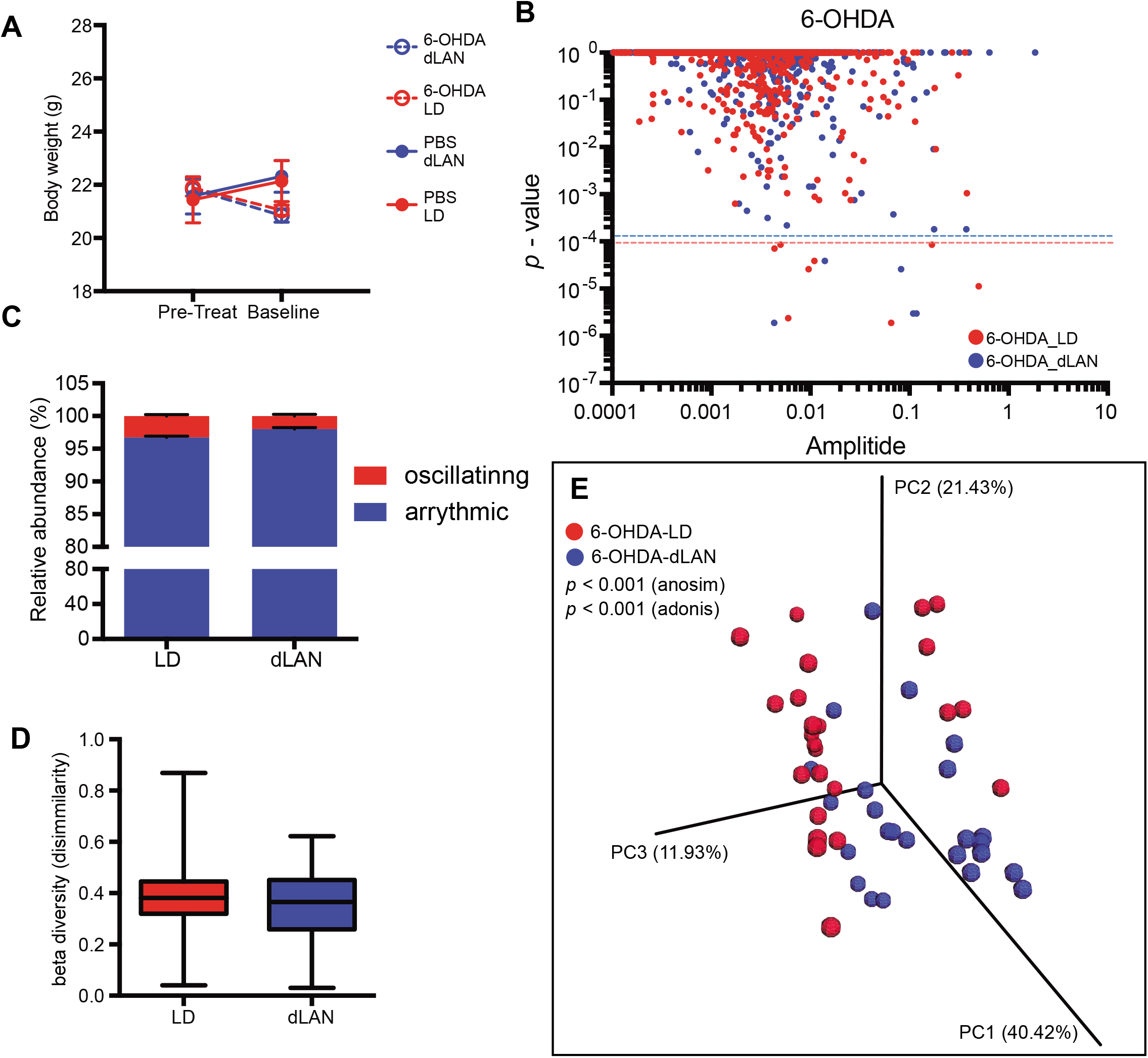
Elimination of sympathetic nerve by 6-OHDA inhibits daily oscillation of gut microbes. (**A**) After 1 week of initial injection (baseline), about 10% of transient body weight loss was observed in 6-OHDA group (open circle) but not in PBS injection group (closed circle) compared to pretreat time point. The body weight recovered after 1 week although 100mg/kg of 6-OHDA was injected every 2 days throughout the experiment. (**B**) OTU based JTK_CYCLE analysis from 6-OHDA treated mice housed under LD and dLAN conditions. Dash lines indicate the Benjamini-Hochberg q-value < 0.05. (**C**) Less than 3% of gut microbes displayed daily oscillations in 6-OHDA injected mice under either LD or dLAN conditions. (**D**) No significant change of gut microbiota beta diversity under dLAN condition in 6-OHDA injected mice. (**E**) PCoA of weighted UniFrac matrix from 6-OHDA injected mice housed under LD or dLAN conditions. Both ANOSIM and Adonis showed significant differences for 6-OHDA treated mice housed under LD or dLAN conditions. n = 4-5 for each condition.

The immune response of intestine cells has been shown to involve in the interaction between host and gut microbes. Therefore, we would like to verify the local molecular marker that participating in light-dependent gut microbe modulation. To see whether dLAN could modulate the gene expression in the intestine, we compared transcriptome data from mice housed under LD or dLAN conditions. We found that 2 weeks of dLAN could influence the gene expression profile in the intestine. Amount 12901 expressing genes, 294 genes are specifically expressed under the dLAN condition and 251 genes are specifically expressed under the control condition (Fig. 6A). For differential expression genes, we found that MMP10 and Iglv3 are upregulated and Nr1d1 is downregulated under the dLAN condition (Fig. 6B). Since the MMP10 gene is involved in immune response and is expressed in the intestine epithelium, we collected intestine samples from mice housed under control and dLAN condition with PBS or 6-OHDA injection similar to previous experimental conditions. Quantitative PCR confirmed that MMP10 is upregulated under the dLAN condition in the intestine. Furthermore, the upregulation can be blocked by the elimination of sympathetic nerve using 6-OHDA (Fig. 6C). Together, these results indicate that environmental light signals transmitted by the ipRGC-sympathetic nerve circuit could modulate the immune response of the digestive tract and is important to drive the daily oscillation of gut microbes. In contrast, ipRGC could influence the composition of gut microbes through the sympathetic nerve independent unidentified pathway (Fig. 6D).

**Figure 6.**
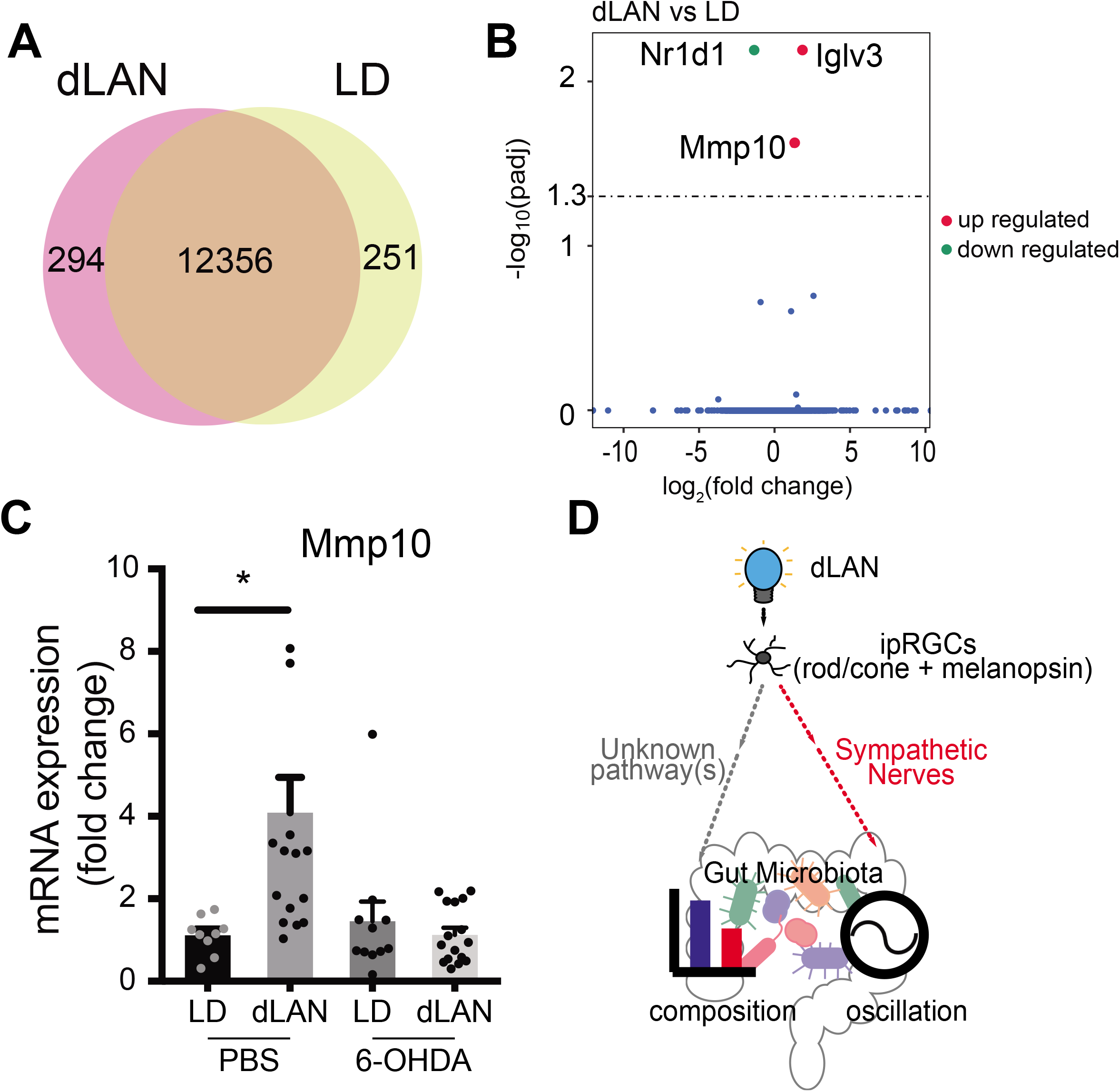
Gene expression profile of intestine under dLAN or LD condition. (**A**) Transcriptome analysis of intestine from mice housed under dLAN or LD conditions. Vann diagram showed that 294 genes are specifically expressed under dLAN condition while 251 genes are specifically expressed under LD condition. (**B**) Nr1d1 gene is significantly down-regulated while MMP10 and Iglv3 are up-regulated in dLAN condition compared to control. (**C**) Quantitative-PCR showed that MMP10 gene expression is up-regulated under dLAN condition. Mice injected with 6-OHDA to eliminate sympathetic nerve do not show a significant difference of MMP10 gene expression between dLAN and LD condition. (**D**) Light-dark cycle information transmitted by ipRGC through sympathetic dependent and independent pathways to influence the daily oscillation of gut microbes and the composition of gut microbiota respectively. n = 4-5 for q-PCR and n = 3 for transcriptome. * p<0.05 with ANOVA.

## Discussion

Our data demonstrate that light input from ipRGCs is a critical regulator of the gut microbiome. We find that gut microbe composition and oscillation are significantly altered in the absence of ipRGC signaling through a sympathetic circuit. Moreover, night time light exposure also alters gut microbe composition and diversity through ipRGCs and non-sympathetic circuit. Together, our results indicate that external light information is an important driving force to shape the gut microbiota in mice through melanopsin and ipRGC circuitry. Since dysbiosis is associated with many metabolic diseases, our results suggest that the light-dark cycle might modulate metabolic status through ipRGCs and gut microbiota interaction.

It has been shown that clock gene(s) are essential for gut microbe daily oscillation since such oscillations are highly attenuated in circadian core gene Per1/2 knockout mice (10). Here we showed that many gut microbes display daily oscillation under normal light-dark cycle similar to previous reports (8, 10, 40). However, oscillations of gut microbes were substantially dampened from mice housed under dLAN, which showed daily circadian rhythm behaviorally and genetically. Furthermore, gut microbe oscillations were highly dampened in the ipRGC-eliminated mice and WT mice housed under constant darkness, both have a free-running endogenous clock. Together, our results suggested that the light-dark cycle is also a strong force to drive the daily oscillation of gut microbes in addition to the circadian clock of the host. Here we hypothesize that distinct species of gut microbe were oscillating in different individuals of mice without ipRGC sending proper light-dark cycle information. Therefore, the discrete oscillating microbes within different mice may be averaged out in JTK cycle analysis. This is similar to the result that peripheral clocks will free run with their own period in vitro without input from the SCN (41). Alternatively, the information of external light-dark conditions could potentially modulate the peripheral clock, which is reported recently (42, 43), to influence the daily oscillation of gut microbe. Consequently, a mismatch of the host circadian clock and external light-dark cycle (e.g. dLAN or constant darkness) disrupted gut microbes, apparently through untimely activation of sympathetic nerves. Another potential hypothesis is that proper entrainment could maintain or magnify the oscillating amplitude of the host circadian clock to provide sufficient driving force for gut microbes. Therefore, the circadian central clock of the host and the light-dark cycle signal could have an additive effect to modulate the daily oscillation of gut microbes through single or multiple neuronal circuitry including sympathetic nerves. Finally, it is possible that an appropriate light input at the correct circadian time is necessary for the daily oscillation of some gut microbes. Thus, synchronization of the external light-dark cycle with the central circadian clock is important to drive the full amplitude of gut microbe daily oscillation. Nevertheless, our conclusion that ipRGC could transmit light-dark cycle information to shape the coherent daily oscillation of gut microbes remains unchanged.

Interestingly, there was a small decrease in the daily oscillation of gut microbes in MKO mice (16%, compared to 20% in WT) exposed to normal LD condition, suggesting that an extrinsic signal from photoreceptor rods and cones through ipRGCs could provide sufficient information to drive daily oscillations of gut microbes. However, dLAN could not significantly alter the overall composition of gut microbiota in MKO mice. This result indicates that melanopsin photo-detection was required to modulate the part composition of gut microbiota. This phenomena of melanopsin discrepancy are similar to the circadian photoentrainment, in which melanopsin is not necessary under certain conditions. Activation of ipRGCs with rods and cones in melanopsin knockout mice is sufficient to entrain animals to the external daily light-dark cycle (44, 45), which is similar to what drives gut microbe oscillation. However, full activation of ipRGC with melanopsin is required for period lengthening under constant light (18, 46, 47), which is similar to systematic modulation of gut microbiota composition. Therefore, information transmitted by ipRGCs to the brain may utilize multiple layers of computation circuits to modulate gut microbiota. Recent studies showed that ipRGC could release both glutamate and GABA (48), or innervates all major types of neurons in the SCN (22). Furthermore, ipRGC input to the SCN and IGL could be involved in the gut microbiota modulation through food entrainment circuitry (49). These complicated circuits may help explain why we observed 2 distinct phenotypes of gut microbiota under dLAN condition.

In addition to daily oscillation modulation, we found that relative abundance of two specific genera *Rikenella* and *Odoribacter* were down-regulated in dLAN condition, while order *RF39*, family *S24-7*, genus *Paraprevotella*, and *Lactobacillus* were up-regulated. It has been shown that *Rikenella* and *Odoribacter* in the gut are associate with non-infective inflammation (50–53). *RF39, S24-7* has been shown to associate with diabetes and obesity (54–56). Although our transcriptome data did not report direct up-regulation of inflammation factors, pathway analysis showed that the immune signaling was different between animal housed under control LD and dLAN condition. Specifically, MMP10 which has been shown to modulate inflammation (57) is up-regulated in mouse housed under dLAN. Therefore, dLAN might be a risk factor for inflammation and metabolic disorders. Indeed, a recent study showed that arrhythmic gut microbiota is linked to patients with type 2 diabetes(48). Since we only house mice under dLAN for 2 weeks, further study could focus on the relationship between chronic dLAN and metabolic or inflammation-related disease. In brief, the brain to gut neuronal circuitry including ipRGC and the sympathetic nervous system could be involved in shaping the gut microbiota through modulating host-microbe interaction molecular pathways in the digestive tract.

In several studies, high-fat diets or environmental stress (e.g. restricted sleep) reduced the richness (alpha diversity) of gut microbiota (7, 8, 10, 40, 58). The high fat diet has been shown to dampen the gut microbe daily oscillation (8). Our gut microbe daily oscillation is indeed slightly lower than previous reports, confirming that a high fat diet could reduce the oscillation amplitude of the gut microbe. Herein, we showed that dLAN further reduced the richness of the gut microbiota under high fat diet, which indicates dLAN is an additional environmental stress for gut microbiota. Finally, the dLAN induced dysbiosis suggests that activation of ipRGC during the night time could disrupt gut microbiota. On the other hand, the gut microbiota composition, alpha, and beta diversity are different between control and MKO- and ipRGC-eliminated mice, suggesting that the presence of melanopsin signaling during the day could also regulate the gut microbiota. Overall, our results together suggest that appropriate light-dark cycle information transmitted by ipRGC is important to maintain a normal gut microbiota.

## Materials and Methods

### Mice

Care of mice and protocols used in this study were reviewed and approved by the Institutional Animal Care and Use Committee at National Taiwan University. Male littermate age-matched controls (*Opn4^Cre/+^*), male melanopsin knockout mice (*Opn4^Cre/Cre^*) and male ipRGC elimination mice (*Opn4^DTA/DTA^*), 8 wk of age, were used. Mice were maintained on a C57BL6/J background.

### Experimental Design

Littermates were housed together under a 12h-12h light-dark (LD) cycle and given a standard diet and water *ad libitum* prior to the experiment. At 8 wk of age, mice were housed individually and randomly assigned to one of two experimental conditions: a normal light/dark cycle (LD) or dim light at night (dLAN). The LD group was exposed to a 12-h light phase, 800 lux and a 12-h dark phase, 0 lux. The dLAN group was exposed to the same 12-h light phase but a 25-lux, 12-h dim light phase. Mice were fed high-fat diets (D12492, 60% fat; Research Diets, Inc., US) during the dark or dim light phase. Mice were either moved to a new cage or feed was removed during the light to limit feeding to the dark or dim light phase. Mice which hid food under bedding were eliminated from the experiment. Daily activity was recorded for 24 h using an infrared camera linked with ANY-maze (Stoelting, Co., US). After 2 wk in experimental light conditions, fecal pellets were collected over 48 h by combining time courses from different mice and using a set period of 24 h (ZT0, ZT4, ZT8, ZT12, ZT16, ZT20) under LD and dLAN conditions for JTK analysis. Fecal pellets were flash frozen (−80 °C) for DNA extraction. For experiments that involved 6-OHDA treatments, 6-OHDA(100 mg/kg, Tocris, UK) was given every 2 d, starting 1 wk prior to the experiments and continuing throughout the experiment. For constant darkness gut microbe oscillation analysis, mice were housed in constant darkness for 2 weeks with temporal restriction feeding between ZT 12 to ZT 24, additional group of constant darkness with ad libitum feeding was also carried out simultaneously. Fecal samples were collected at ZT0, ZT4, ZT8, ZT12, ZT16, ZT20 across 48-hour period for LD and dLAN group. For constant darkness and Opn4^DTA/DTA^ groups, after sample collection, mice were transferred to wheel running cage and the onset of individual mouse was calculated using ClockLab (Actimetrics, US). The specific collection time was matching to the calculated circadian time for analysis.

### Metagenomic library preparation and 16S sequencing

The DNA was extracted from ~25 mg of feces, using a QIAamp DNA Mini Kit (Qiagen, Germany) according to the manufacturer’s instructions and 40 ng of DNA was used in PCR amplification and 16S rRNA gene sequencing. Amplicons spanning the variable region 4/5 (V4/5) of 16S rRNA gene were generated using the following primers:

5’-TCGTCGGCAGCGTCAGATGTGTATAAGAGACAGAYTGGGYDTAAAGNG-3’ and 5’-GTCTCGTGGGCTCGGAGATGTGTATAAGAGACAGCCGTCAATTYYTTTRAGTTT-3’.

The PCR products were cleaned and barcoded using Nextera^®^ Index Kit (Illumina Inc, US). Final products were purified, concentrated and DNA quality verified. The library was sequenced on an Illumina MiSeq platform, with 250 bp paired-end sequencing.

### Microbiota sequence analysis

All analysis is following the procedure of Thaiss et. al., 2014 *Cell* and Zarrinpar et. al., 2014 *Cell metabolism*. Reads were classified with QIIME (Quantitative Insight into Microbial Ecology) 1.90. In brief, operational taxonomic units (OTU) were picked using close reference methods at 97% sequence similarity against the Greengenes Database (The Greengenes Database Consortium, Version 13.8), taxonomies were assigned using the uclust consensus taxonomy assigner, and an OTU table was created. Samples with < 2,000 reads were discarded and removed from further analysis. The average number of reads for the samples was 38,361. For beta diversity analysis, unweighted and weighted UniFrac distances were calculated, and principal coordinate analysis plots were generated. Dissimilarity matrixes were also used to compare in-group differences between microbe samples. To determine the diurnal fluctuations of each OTU, the percentage of reads was calculated for each sample and averaged for each time point and data analyzed using the JTK_CYCLE program. The significances of cyclic OTUs were achieved by both permutation-based adjusted p values and Benjamini-Hochberg q-values were < 0.05. Linear discriminant analysis effect size (LEfSe) was used to detect significant changes in the relative abundance of microbial taxa between groups via online Galaxy Browser. Significant thresholds were set as default settings: alpha < 0.05 for a factorial Kruskal-Wallis test among classes and LDA > 2.0 for logarithmic LDA score.

### Quantification of circadian gene expression

Livers were homogenized using a tissue homogenizer (Minilys lyser, Bertin Corp., France) and micro-dissected SCN tissues were lysed in lysis buffer by repeated aspiration through a syringe. Total RNAs were then extracted using Quick RNA Mini Prep (Zymo, R1054, US) according to the manufacturer’s instructions. Then, cDNAs were synthesized from total RNA extracts using iScript cDNA Synthesis Kit (Bio-Rad, US) and cDNA products stored at 4 °C before use. Thereafter, qPCR was performed using iQ SYBR Green Supermix QPCR kit (Bio-Rad) and detected in a CFX96 thermal cycler and detection system (Bio-Rad). Primers for circadian gene detection are listed below. Circadian genes mRNA expressions were normalized by 18S ribosomal RNA (internal control).

Primer list for QPCR

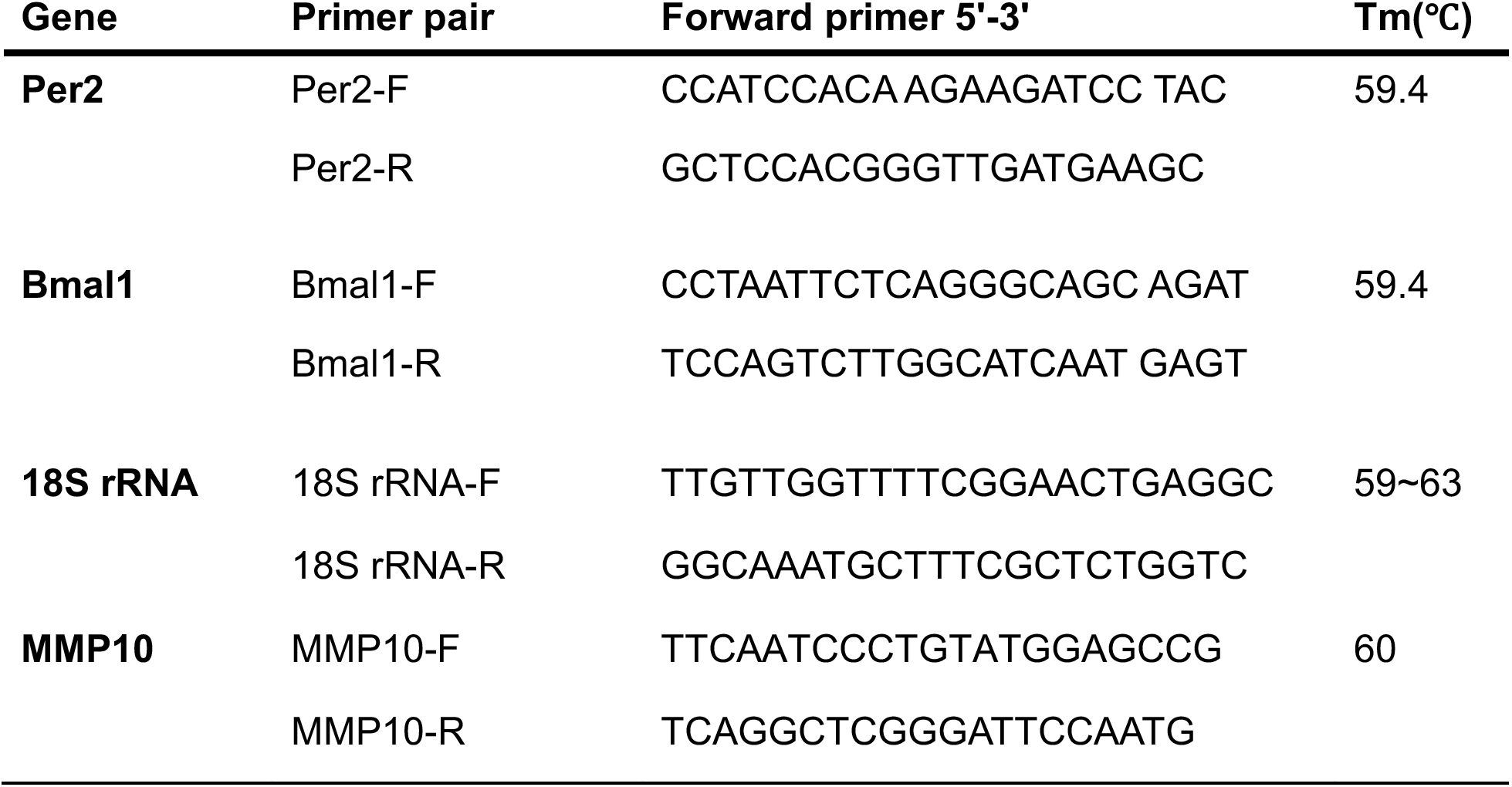

### Immunostaining

For immunohistochemistry staining, the mice were perfused with PBS for 2 min and 4% paraformaldehyde (PFA) for 15 min, brains were removed and post-fixed in 4% PFA for 2 hours. After fixation, mouse brains were sectioned coronally with 80 μm thickness using a vibratome. Brian slices were first incubated with 3% H_2_O_2_ in PBS for 10 min, then blocked in 5% goat serum, 1% BSA, and 0.2% Triton-100 in PBS (0.1M) for 2 hours at room temperature. Subsequently, the brain slices were incubated overnight at 4°C with a blocking solution containing Rabbit anti-Per2 (1:1000; Alpha Diagnostic, PER21-A) antibodies. After washing in PBS (0.1M), the brain was incubated with VECTASTAIN ABC KITS (PEROXIDASE, Vector Lab) following manufactory instruction. The DAB staining performed using DAB Peroxidase (HRP) Substrate Kit (Vector Lab) following manufactory instruction and then mounted with VECTASHIELD^®^ (Vector Lab) for imaging. Bright-field images were collected using a Zeiss Z1 microscope with a 20X objective.

### Transcriptome

Mice aged 8 weeks were randomly separated into two groups and housed individually under different light conditions: normal LD cycle (light 800 lux; dark 0 lux) and group in dim light at night (light 800 lux; dim light 40lux) for two weeks. All mice access to ad libitum filtered water and restricted high-fat diet (HFD) (D12492, 60% kcal of fat) at night time. Mice were transferred to day time cage and night time cage to ensure the food residue at night would not be taken at day time. We measured mice body weight and food intake every week. Mice were anesthetized with avertin (2, 2, 2-Tribromoethanol) at ZT 14. Small intestine tissue was moved out and dissected on ice divided into three sections depending on length. The intestinal contents were cleaned up by PBS and the small intestine was frozen immediately by liquid nitrogen. RNA was extracted from intestine tissue by RNA kit (Zymo Quick-RNA miniprep, R1055) for transcriptome sequencing.

### Statistical analyses

Unless stated, all statistical analyses were performed using the Prism6 software. To compare differences between time points and light conditions, data were analyzed by two-way ANOVA, followed by multiple comparisons with Bonferroni’s *post hoc* test. For comparing data between two groups, an unpaired Student’s *t*-test was applied. In all cases, differences between groups were considered statistically significant at p < .05.

## Acknowledgments

This work was supported by the Taiwan Ministry of Science and Technology grant MOST 103-2628-B-002-001-MY3 (to S.-K.C.). We thank the Technology Commons, College of Life Science at National Taiwan University for technical assistance with NGS, Dr. Yi-Juang Chern and Dr. Bon-Chu Chung for their support and discussions, and Dr. Allan Y.-C. Chang for writing assistance. We also thank Dr. Samer Hattar for providing transgenic mice for this study. The authors declare no competing financial conflicts of interest.

## Author contributions

H-T. Y., S-K. C. and C-C. L. designed the experiment and wrote the manuscript. C-C. L., T-H. L., Y-F. Z. and I-C. L. performed all experiments and analyses.

