## supplemental text and figures for "External light dark cycle shapes gut microbiota through intrinsically photosensitive retinal ganglion cells"

### Supplementary information text

*Mice housed under dLAN condition showed similar circadian photo-entrainment to normal LD condition.*

The light intensity for dLAN during the subjective night is 25 lux (around $2.7*10^{12}$ photon/s/$cm^2$ at 400-500 nm wavelength range), which is above the threshold for all characterized photoreceptors in the retina including rods, cones, and ipRGCs. Since temporal restriction feeding during the subjective night time could block dLAN induced circadian clock desynchronization [1], and mice with timed high fat diet exhibit less stressed phenotype compared to mice with timed low-fat diet [2], we performed temporal restriction of high fat diet (Supplementary Fig. 1A) to minimize circadian disruption during the experimental period [3]. After 2 weeks, we first confirmed that the daily activity rhythms were entrained to 24 hours cycle from mice housed under LD or dLAN conditions (Supplementary Fig. 1B). In addition, the expression patterns for clock genes *Bmal1* and *Per2* in the SCN and liver were similar between mice housed under dLAN and LD condition, although the Bmal1 is slightly dampened in the SCN while the peak time of Per2 is advanced only in the liver (Supplementary Fig. 1C). The Per2 protein expression patterns in the SCN were also similar between dLAN and LD condition (Supplementary Fig. 1D and 1E). Together, our results indicated that the central clock remains entrained and show daily oscillation similar to the normal LD cycle when mice were exposed to 25 lux of light during the subjective night time under temporal restricted high fat diet condition.

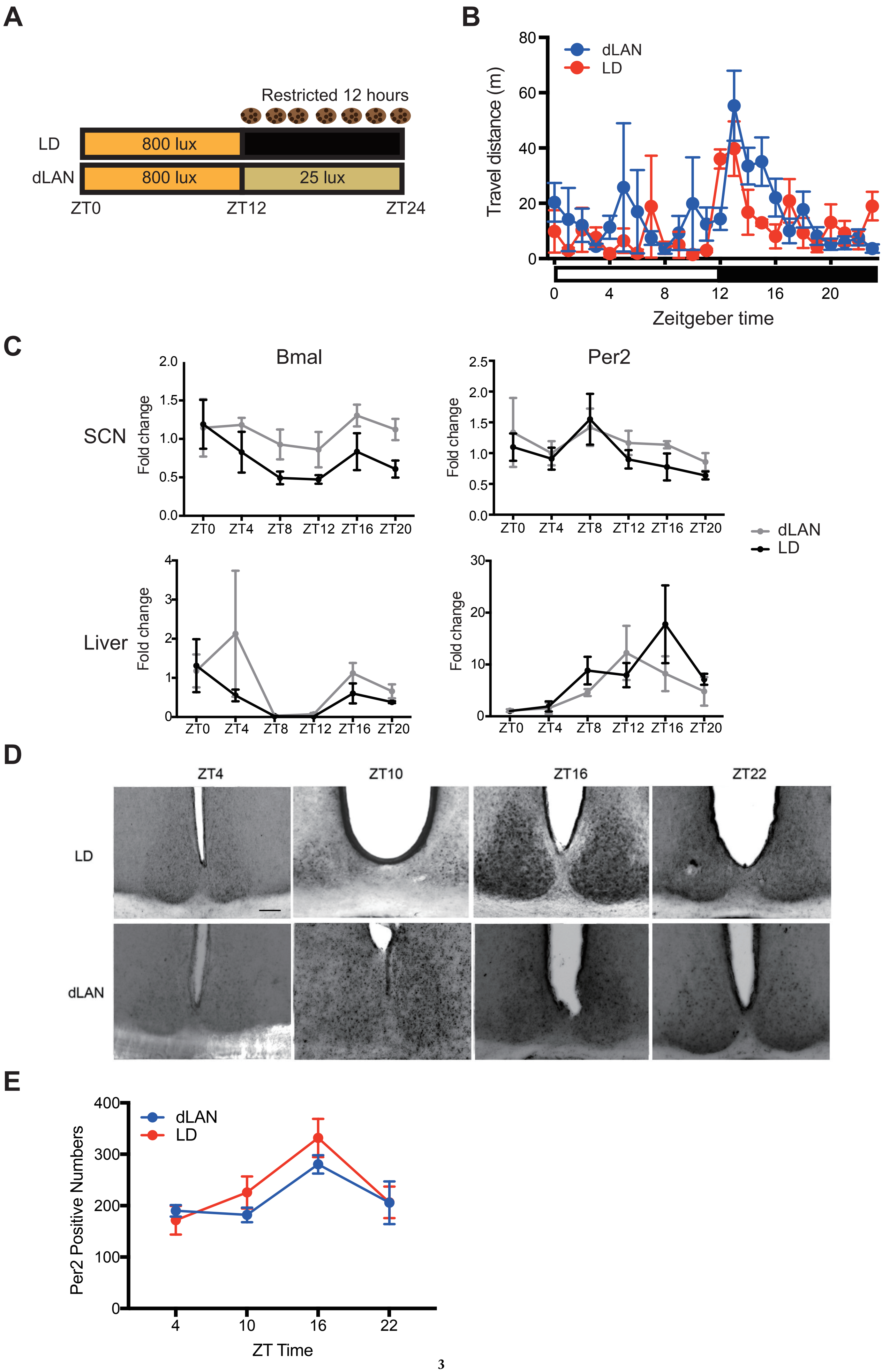




Supplementary Fig. 1

**Figure S1. Mice housed in dLAN condition display similar activity and clock gene expression patterns to LD condition**

(**A**) Schematic representation of the experimental design for mice exposed to dim light at night (dLAN) or normal light/dark (LD). LD mice were housed under a 12h-12h LD cycle with 0 lux during the dark phase; dLAN mice were housed under 25 lux during the dark phase. Food availability was restricted to the dark phase for both conditions. (**B**) Temporal activity profile of mice, plotted by the moving distance in the home cages per hour, under dim light at night (dLAN, red) or normal light/dark (LD, blue) conditions. n = 5 for each group. (**C**) QPCR analysis of expression levels of the circadian clock genes *Bmal1* and *Per2* in the suprachiasmatic nucleus (SCN) and liver from Control mice housed under conditions of dLAN (gray) and LD (black). **(D)** Representative images of Per2 immuno-positive cells in the SCN at ZT2, ZT8, ZT14, and ZT20 from control mice housed under LD (upper) and dLAN (lower). (**E**) Quantification of Per2 immuno-positive cell in the SCN. There is no significant difference in the Per2 positive cell number between mice housed under LD and dLAN condition. n = 4-5 for qPCR, n = 3 for IHC. Scale bar is 100 μm for (D). Data are presented as mean ± SEM.

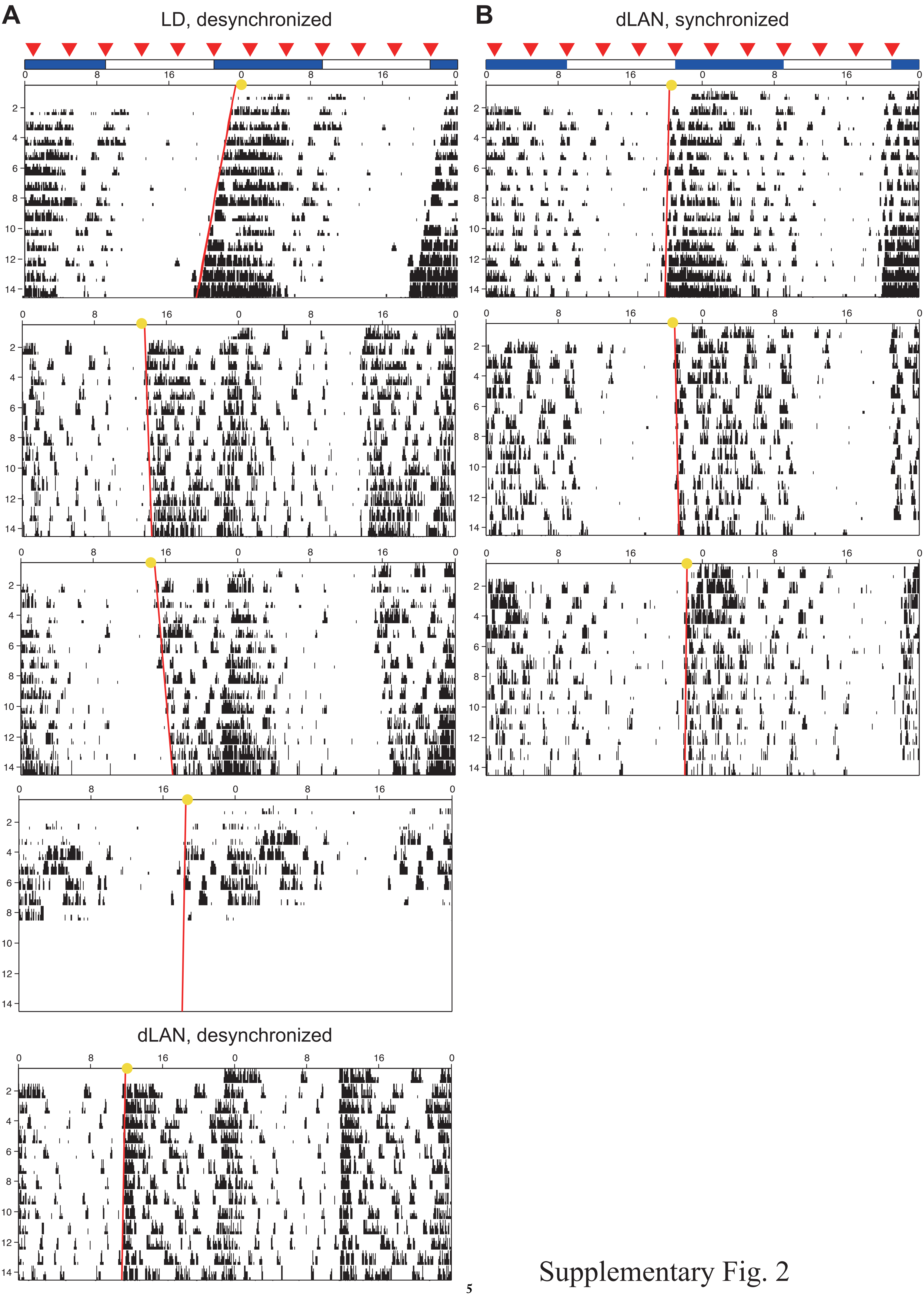


Supplementary Fig. 2

**Figure S2. Actogram of Opn4$^{DTA/DTA}$ mice**

Wheel running activity of Opn4$^{DTA/DTA}$ mice immediately after final fecal sample collection. (**A**) Opn4$^{DTA/DTA}$ mice, which activity onsets are at least 3 hours in advance or delay to the light off, housed under LD or dLAN condition. (**B**) Opn4$^{DTA/DTA}$ mice, which activity onsets are advance or delay to the light off less than 3 hours, housed under LD or dLAN condition. Yellow circles indicate predicted onset for fecal sample collecting days. Red triangles indicate fecal sample collecting times.
